# Apelin-VEGF-C mRNA delivery as therapeutic for the treatment of secondary lymphedema

**DOI:** 10.1101/2023.01.05.522869

**Authors:** A. Lamaa, J. Creff, E. Benuzzi, f. Pujol, T. Draia-Nicolau, M. Nougué, L. Verdu, F. Morfoisse, E. Lacazette, P. Valet, B. Chaput, F. Gross, R. Gayon, P. Bouillé, J. Malloizel-Delaunay, A. Bura-Rivière, A.C. Prats, B. Garmy-Susini

## Abstract

Secondary lymphedema (LD) corresponds to a severe lymphatic dysfunction leading to the accumulation of fluid and fibrotic adipose tissue in a limb. Here, we identified apelin (APLN) as a powerful molecule for regenerating lymphatic function in LD. We identified the loss of APLN expression in lymphedematous arm compared to normal arm in patients. The role of APLN in LD was confirmed in APLN-knockout mice, in which LD is increased and associated with fibrosis and dermal backflow. This was reversed by intradermal injection of APLN-lentivectors. Mechanistically, APLN stimulates lymphatic endothelial cell gene expression and induces the binding of E2F8 transcription factor to the promoter of CCBE1 that controls VEGF-C processing. In addition, APLN induces Akt and eNOS pathways to stimulate lymphatic collector pumping. Our results show that APLN represents a novel partner for VEGF-C to restore lymphatic function in both initial and collecting vessels. As LD appears after cancer treatment, we validated the APLN-VEGF-C combination using a novel class of safe and non-integrative RNA-delivery LentiFlash^®^ vector that will be evaluated for phase I/IIa clinical trial.

## INTRODUCTION

Lymphedema (LD) is a multifactorial condition that substantially affects the quality of life of patients (Greene *et al*, 2012; Hoffner *et al*, 2018). It is characterized by a failure of the lymph transport back to the blood circulation due to a lymphatic dysfunction that occurs after genetic mutation (primary LD) or arises after cancer treatments (secondary LD)(Mortimer & Rockson, 2014). The major hallmark of LD is the development of fibrosis into the skin and the adipose tissue (AT) as lymphostatic fibrosis defines the stage of LD from reversible to elephantiasis stages (Mortimer & Rockson, 2014). Once fibrosis develops, tissues become denser leading to a lymphatic vessel obstruction that worsens LD. Importantly, fibrosis also affects collecting lymphatic pumping and increases limb swelling (Baik *et al*, 2022; Kataru *et al*, 2019). A large number of cytokines and peptides are known to selectively improve adipocyte metabolism, endothelial function or tissue fibrosis. However, the bioactive peptide apelin (APLN) combines these beneficial effects as a whole. APLN is the endogenous ligand of the G-protein-coupled receptor APJ, and is a critical actor of the fibrosis protection in many organs including heart, lung and kidney (Huang *et al*, 2016; Pope *et al*, 2012). The first evidences of the link between APLN and lymphatic vasculature were identified in a tumoral environment as APLN stimulates both hemangio- and lymphangiogenesis (Berta *et al*, 2014). In particular, APLN stimulates NO production via PI3K/Akt signalling in blood endothelial cells (Busch *et al*, 2015). Also, our group found that APLN was able to restore the lymphatic shape of precollecting vessels after cardiac ischemia suggesting that APLN may represent a good candidate to restore the lymphatic shape in injured tissues (Tatin *et al*, 2017).

Based on data accumulated throughout 20 years of clinical trials, gene therapy has an acceptable safety profile for vascular diseases (Gupta *et al*, 2009). However, more rigorous phase II and phase III clinical trials have failed to demonstrate that growth factors administered as single-agent gene therapies are beneficial in the patients with cardiovascular diseases. Also, secondary LD related to cancer treatment is an ongoing challenge, which forces therapies to avoid any effect on cancer recurrence. Recent Covid19 vaccines has demonstrated that mRNA can be used as a way of delivering therapeutic proteins. But compared to vaccines, mRNA therapeutics requires a 300-4,000-fold-higher level of protein expression to reach a therapeutic effect when they are delivered through lipid nanoparticles due to a poor entry efficiency into target cells and synthetic RNA stability (Rybakova *et al*, 2019). To overcome these limits and enhancing both duration and protein expression level *in vivo*, we propose to use a biological RNA delivery technology LentiFlash^®^ to transfer 2 different mRNAs into lymphedema tissues. Therefore, we propose to combine APLN to VEGF-C, the major lymphangiogenic factor and the only one molecule that has been evaluated in clinical trial for secondary LD treatment (Hartiala *et al*, 2020). VEGF-C biological effect is enhanced by the collagen- and calcium-binding EGF domains 1 (CCBE1) along with a disintegrin and metalloproteinase with thrombospondin motifs-3 (ADAMTS3) protease (Jha *et al*, 2017). CCBE1 is a secreted molecule involved in lymphatic vasculature development and mutations in *CCBE1* were identified in patients with Hennekam syndrome, a rare autosomal recessive disorder of lymphatic development leading to primary LD (Hogan *et al*, 2009; Van Balkom *et al*, 2002).

Here, we found a significant decrease of APLN in biopsies from patients who develop LD after breast cancer. We showed that APLN stimulated gene expression in LEC through E2F8, a part of E2F family of transcription factors that is an important regulator of lymphangiogenesis in zebrafish and mice after binding to CCBE1 and Flt4 promoters (Lammens *et al*, 2009; Logan *et al*, 2005; Weijts *et al*, 2013). We identified APLN as a crucial player in the NO-induced lymphatic pumping by stimulating eNOS phosphorylation in LEC. Thus, we performed a multiple gene therapy by combining VEGFC with APLN to obtain a synergistic effect necessary to restore the lymphatic function. Our study provided evidences for the use of APLN-VEGF-C combination in patients who developed LD after cancer treatments. Here, we present the preclinical study using APLN-VEGFC mRNA delivery vectors that will be used in a Phase I/II gene therapy clinical trial for the treatment of patients who developed LD after breast cancer.

## RESULTS

### LD exhibits increased lymphatic capillary diameter and poor collecting drainage

Secondary LD occurs months, sometimes years after cancer treatment suggesting that this pathology is not only a side effect of the surgery. Lymphofluoroscopy of woman who developed LD after breast cancer showed hypervascularized dermis with tortuous lymphatic capillaries associated with a strong desmoplastic reaction and dermal backflow (Fig. 1A). Lymphoscintigraphy is the primary imaging modality used to assess lymphatic system dysfunction. It has been considered the criterion standard for decades (Munn & Padera, 2014; Szuba *et al*, 2003). Lymphoscintigraphy revealed a severe decrease in lymphatic collecting vessels detection and axillary lymph node perfusion after radiotracer injection (Fig. 1B). Histological analysis showed an increase in dilated overloaded lymphatic vessels in the skin (Fig.1C-E). This was associated with a strong fibrosis development in both skin (Fig.1F) and AT (Fig.1G). Surprisingly, no major difference in genes involved in lymphangiogenic factor maturation was observed except for CCBE1 that was significantly downregulated in LD (Fig.1H). As LD is characterized by a strong accumulation of fibrotic AT in the limb, we also evaluated adipokines expression in the lymphedematous AT compared to the normal arm (Fig. 1I). We found a significant decrease in APLN expression in LD whereas no difference was observed for other adipokines (Fig.1I).

**Figure 1:**
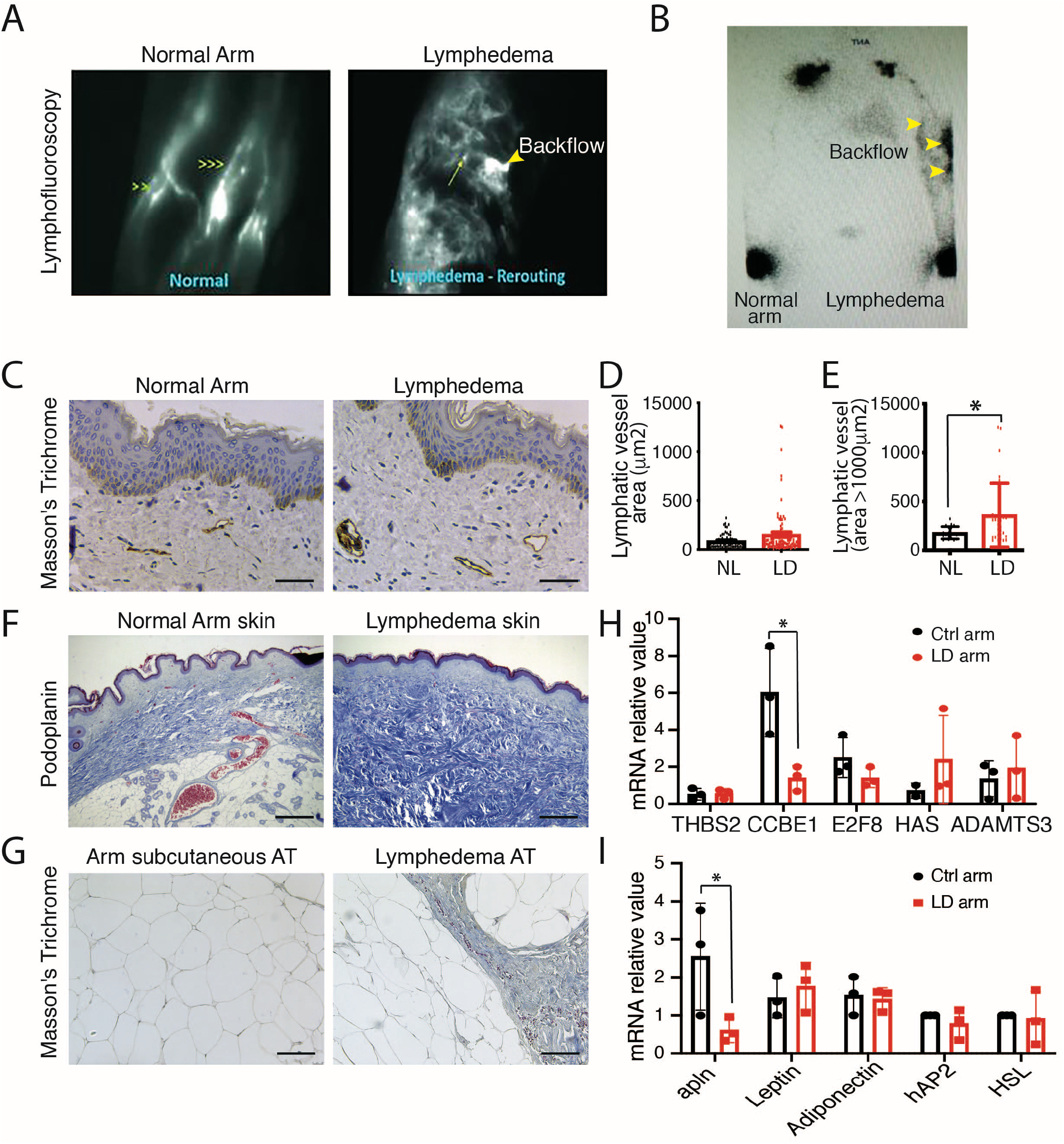
Reduced APLN expression in human LD. **A.** Lymphofluoroscopy of the upper limb LD shows dermal lymph backflow associated with hypervascularization of tortuous initial lymphatics (right panel) compared to normal arm (left panel). **B.** Lymphoscintigraphy of woman who developed LD after breast cancer shows reduced but lasting lymphatic drainage and lymph node perfusion**. C.** Immunodetection of the lymphatic networks in the LD skin shows overloaded dilated lymphatic vessels (scale bar: 50 μm). **D.** Quantification of the lymphatic vessel diameters (*p<0.05). **E.** Quantification of the dermis lymphatic density (p<0.05). **F.** Masson’s trichome staining of the human Lymphedematous skin shows strong fibrosis. **G.** Masson’s trichome staining of the human LD subcutaneous AT shows fibrosis. **H.** Quantitative RT-PCR analysis of the genes involved in fibrosis and VEGFC maturation in dermolipectomies from patients with LD. **I.** Quantitative RT-PCR analysis of the adipokines in dermolipectomies from patients with LD.

### Impaired lymphatic healing in APLN knockout mice

APLN was previously described by our group to improve lymphatic vessel normalization in heart after cardiac ischemia (Tatin *et al*., 2017). To investigate the role of APLN in secondary LD, we used a LD mouse model previously developed in the laboratory (Morfoisse *et al*, 2018). We performed second mammary gland mastectomy associated with axillary and brachial lymphadenectomy on the upper left limb in APLN-KO mice (Fig. 2A). Using this model, reproducible LD developed after 2 weeks to progressively return to normal after 4 weeks. In APLN-KO mice, LD was significantly maintained after 4 weeks showing the failure to restore the lymphatic function (Fig. 2A). Lymphatic capillaries were next investigated using lymphangiography after footpad injection of FITC-Dextran (Fig. 2B). We observed a strong dermal backflow in APLN-KO mice (Fig. 2B). Histological analysis of the lymphatic showed no difference in lymphatic basal density between WT and APLN-KO mice (Fig. 2C and D). In contrast, in LD, the skin lymphangiogenesis was significantly reduced in APLN-KO mice compared to WT mice (Fig. 2C and D). This was associated with an increase in skin fibrosis in both WT and APLN-KO mice as shown using Masson’s trichrome staining (Fig. 2E and F).

**Figure 2:**
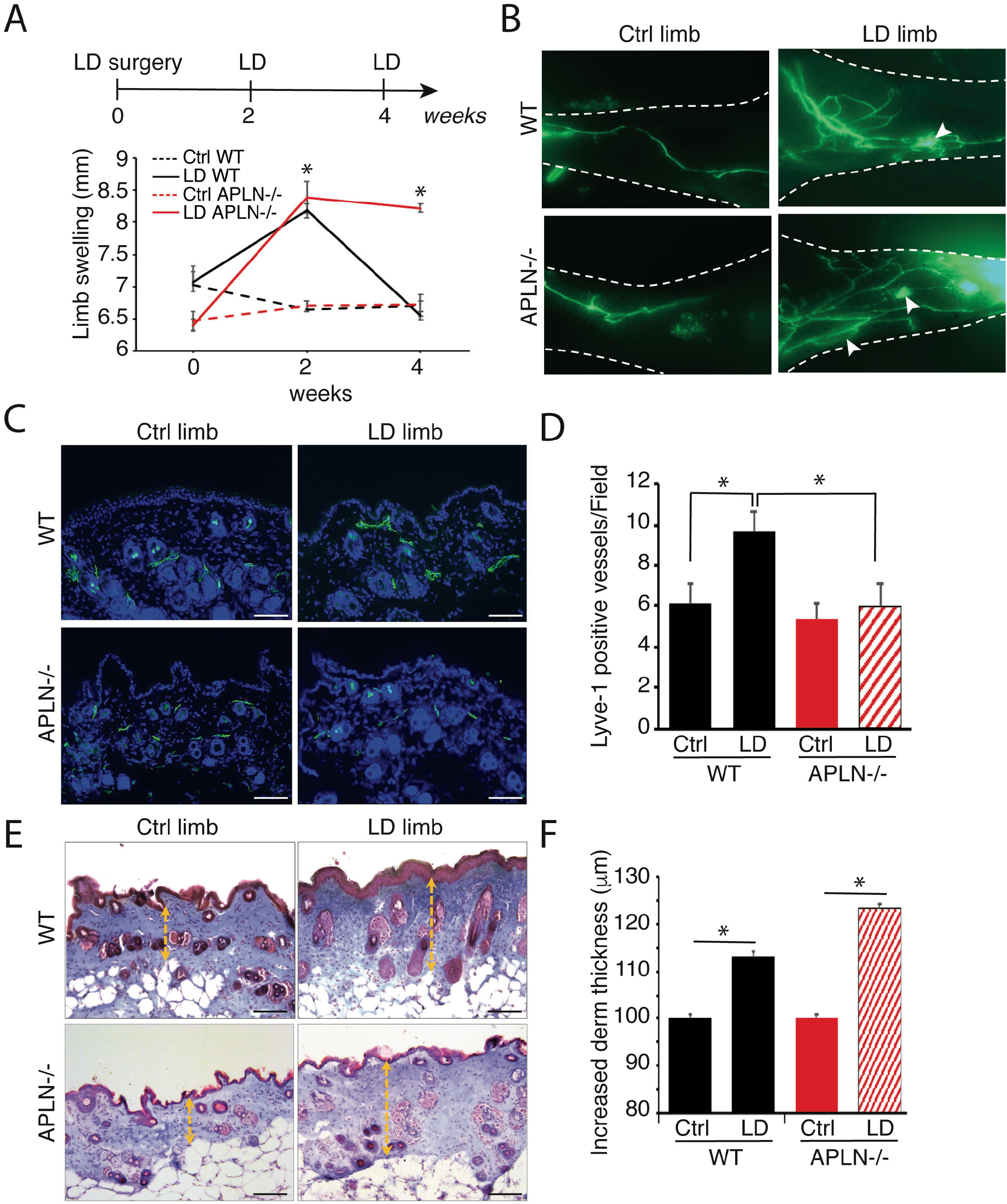
LD increases in APLN-KO mice. **A.** Schematic of the experimental design of secondary LD mice model. Quantification of proximal limb swelling 2 and 4 weeks after surgery on control limb, LD limb from APLN KO mice and control littermates (n=9). **B.** Lymphangiography reveals dermal backflow (white arrows) and pathological remodeling of lymphatic vessel after LD in APLN LO mice. **C.** Lyve-1 immunodetection of the skin lymphangiogenesis in APLN KO mice (scale bar: 50 μm). **D.** Quantification of lymphangiogenesis in the skin from APLN KO mice (*p<0.05). **E.** Masson’s trichrome staining of LD in APLN KO mice (scale bar: 50 μm). **F.** Quantification of dermis fibrosis in APLN KO mice (*p<0.05).

### APLN possesses regenerative function on lymphatic vessels in LD

To evaluate the effect of APLN on lymphatic healing, mice received an intradermal injection of APLN-expressing lentivector (LV-APLN) in the lymphedematous limb. Remarkably LD was significantly reduced in APLN-treated mice (Fig. 3A). Level of circulating APLN was verified by ELISA dosage on mouse plasma showing and increase in plasmatic APLN concentration in LV-APLN-treated mice (Fig. 3B). Lymphatic collecting drainage was next investigated using lymphangiography (Fig. 3C). We observed that secondary LD induced a pathological remodelling of lymphatic vessels with disorganized and abnormal vessel morphology and increased number of branching regarding the control limb. Lymphatic leakage (dermal backflow) was also observed revealing a dysfunction in superficial overloaded capillary network due to the lack of deeper collector pumping (Fig. 3C). On the contrary, APLN-treated mice displayed an improvement in the lymphatic shape with normalized morphology and decreased number of vessels branching. Importantly, we did not observe dermal backflow in APLN-treated mice suggesting an improvement of lymphatic function (Fig. 3C). Using Masson’s trichrome coloration, we observed an increase of the dermis thickness in LD limb consistent with the development of fibrosis (Fig. 3D). In APLN-treated mice, we did not observe any thickening of the dermis (Fig. 3D and E) reflecting an improvement of LD pathology. Interestingly, we observed an increase in circulating VEGF-C, the major lymphangiogenic factor, after LV-APLN treatment suggesting that APLN may in part regulate VEGF-C protein synthesis (Fig 3F). Positive control was performed using VEGF-C-expressing lentivector (LV-VEGFC)(Fig. 3F). The number of blood vessels was assessed using CD31 immunostaining (Fig. EV1). As expected, we did not find changes in the number of CD31-positive vessels in this model of LD (Fig. EV1)(Morfoisse *et al*., 2018). However, as previously described in the literature, treatment with APLN lentivector led to an increase of angiogenesis (Fig. EV1) (Wysocka *et al*, 2018). In parallel, lymphangiogenesis was evaluated using Lyve1 immunostaining on skin sections (Fig. 3G-I). In line with lymphangiography results, an increased number of Lyve1-positive vessels was observed in LD limb in comparison to control limb without significant difference when comparing control to LV-APLN treated mice (Fig. 3H). In contrast, we found that APLN promoted significant dilatation of lymphatic vessel (Fig. 3I). Overall, our results indicate that APLN has beneficial role on secondary LD by acting on lymphatic vessel plasticity and dilatation.

**Figure 3:**
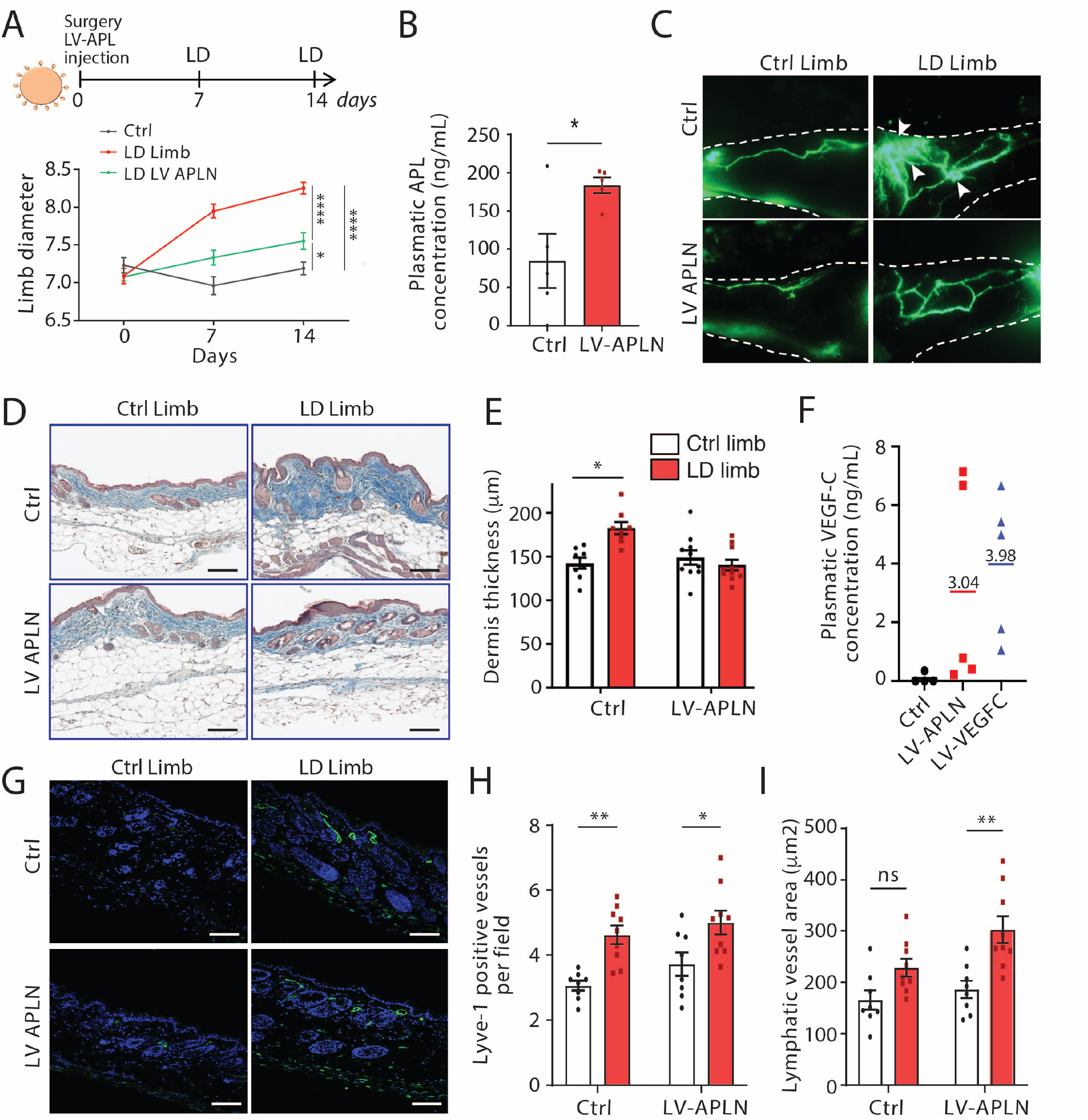
APLN prevents secondary LD. **A**. Schematic of the experimental design of secondary LD model in mice injected APLN lentivector (LV-APLN). Quantification of proximal limb swelling at 7 and 14 days after surgery on control limb, LD limb (n=17) or LD treated with APLN lentivector (n=20)(*p<0.05). **B.** EIA dosage of circulating APLN in plasma of control (n=5) or APLN treated mice (n=5). **C.** Lymphangiography reveals pathological remodeling of lymphatic vessels and dermal backflow in LD that is reversed by LV-APLN (n=10). **D.** Masson’s trichrome staining of the skin from mice with LD treated or not with APLN (scale bar: 50 μm). **E.** Quantification of dermis thickness (*p<0.05). **F.** EIA dosage of circulating VEGF-C in plasma of control (n=5) or APLN-treated mice (n=5). **G.** Lyve-1 immunodetection of the skin lymphangiogenesis in APLN-treated mice (scale bar: 50 μm). **H.** Quantification of lymphangiogenesis in APLN-treated mice (*p<0.05, **p < 0.01). **I.** Quantification of lymphatic dilatation in APLN-treated mice (**p < 0.01).

### APLN controls LEC gene expression

In order to investigate molecular mechanisms regulated by APLN in LEC, we performed a global transcriptomic analysis on LEC stimulated 24 hrs with conditioned media containing APLN or conditioned media obtained from control NIH3T3 (Fig. 4A-H). Differential DESeq analysis revealed that 217 genes were deregulated (p.adj<0.05 and Log2 fold change < −0.5 or >0.5) with 94 genes up regulated and 123 genes down-regulated (Fig. 4A). Top 30 down- or up-regulated are displayed on heat maps (Fig. 4B) and complete lists are given in Table S1 and Table S2. Gene ontology (GO) analysis of down-regulated genes revealed that no biological process is significantly affected in APLN treated HDLEC. In contrast, GO analysis for biological process revealed that up-regulated genes were enriched (FDR < 0.05) for terms related to extracellular matrix (ECM) remodelling and signalisation (Fig. 4C) including COL1A, FBN, ADAMTS2, and CCBE1 (Fig. 4B and C). However, most of this gene induction was not validated by RT-qPCR on HDLEC (Fig. 4D-F) except for CCBE1 whose induction was strongly confirmed (Fig. 4G). CCBE1 protein is required for the activation of VEGF-C along with the ADAMTS3 protease by enhancing the cleavage activity of ADAMTS3 and by facilitating the maturation of VEGF-C into its bioactive form. Interestingly, APLN also stimulated the expression of E2F8, the CCBE1 transcription factor (Fig. 4H). We then postulated that APLN could participate to VEGF-C maturation by increasing E2F8 DNA binding on CCBE1 promoter. To answer this question, we performed chromatin immunoprecipitation (ChIP) of E2F8 in APLN-overexpressing HDLEC (Fig. 4I-K). We found that APLN significantly increased E2F8 binding to CCBE1 promoter (Fig. 4J). Interestingly, APLN also induced E2F8 binding to E2F1 transcription factor promoter suggesting a role in other biological functions (Fig. 4K) (Wells *et al*, 2002). Altogether, these data indicated that APLN regulates HDLEC gene expression.

**Figure 4:**
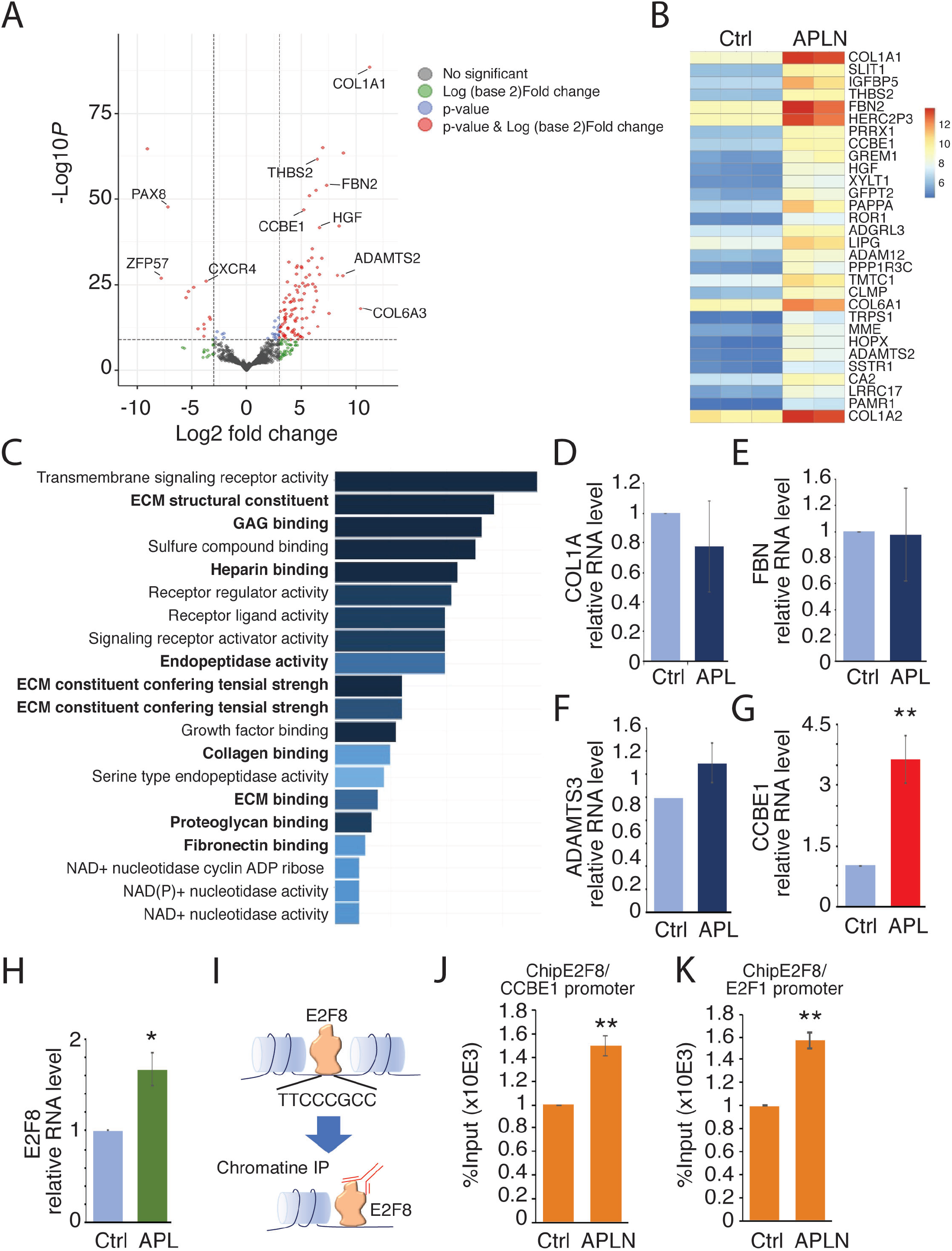
APLN controls lymphatic endothelial cell gene expression. **A.** Bulk RNA-sequencing in HDLEC treated with APLN-conditioned medium. Volcano plot showing log2FC (Fold change) values calculated between control and APLN-treated HDLECs for 24h. Red and blue dots: significantly (*p value adjusted* < 0.05) up- (log2FC > 0.5) and downregulated genes (log2FC < −0.5) respectively. **B.** Heat map of the top 30 significantly up regulated genes (p value adjusted <0.05 and log2FC > 0.5). Expression levels are plotted as log10 normalized counts for each sample. Red represents higher FC; Dark blue represents lower FC. **C.** Top significantly (FDR < 0.05) enriched Gene Ontology (GO) terms for biological processes of significantly upregulated genes after APLN treatment at 24h time point. **D-H.** qRT-PCR validation of COL1A1 (D), FBN (E), ADAMTS3 (F), CCBE1 (G) and E2F8 (H) in APLN-stimulated HDLEC (**p<0.05). **I.** Schematic representation of chromatin immunoprecipitation (Chip) by E2F8. **J.** Chip analysis of E2F8 on CCBE1 promoter (**p < 0.05). **K.** Chip analysis of E2F8 on E2F1 promoter (**p < 0.05).

### APLN stimulates LEC function through Akt/eNOS signalling

Next, we investigated the effect of APLN on HDLEC at the cellular level. APLN is known to activate Erk and Akt signalling *in vitro* in human dermal LEC (HDLEC) (Berta *et al*., 2014; Kim *et al*, 2014). In line with the vasodilation phenotype (Fig. 3), we postulated that APLN beneficial effect on LD is in part mediated by AKT/eNOS pathway. To this end, we stimulated HDLEC with conditioned media obtained from LV-APLN transduced NIH3T3 previously depleted for VEGF-C. APLN synthesis was validated by RT-qPCR on NIH3T3 (Fig. 5A) and by ELISA (Fig. 5B). Stimulation of HDLEC by conditioned media was confirmed by evaluating AKT et ERK pathway during 24 hrs time course. Medium containing VEGF-C was used as positive control (Fig. 5C). HDLEC responded to both VEGF-C and APLN after 30 min as we observed a strong activation of AKT and ERK (Fig. 5C and D). This was associated with an activation of HDLEC migration by APLN (Fig. 5E and F) and an improvement of the actin cytoskeleton remodeling showing cortical actin rim compared to stress fibers observed in negative control (Fig.5G). However, no effect was observed on cell junction (Fig. 5G). Importantly, eNOS phosphorylation was observed in HDLEC in response to APLN and to VEGF-C, suggesting that both APLN and VEGF-C stimulate lymphatic dilatation through eNOS pathway (Fig. 5H and I).

**Figure 5:**
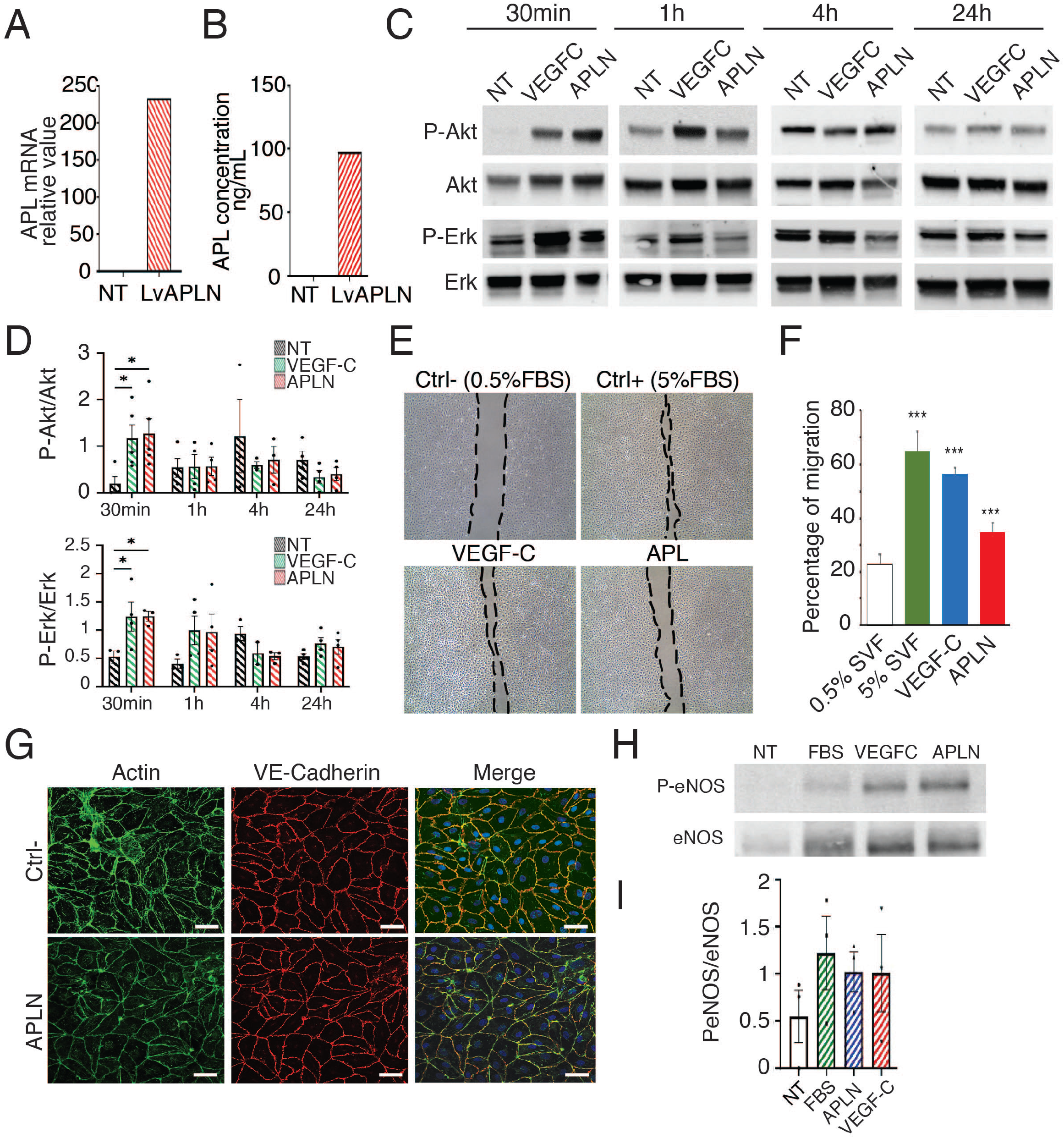
APLN plays a role in LD through Akt/eNOS activation in lymphatic endothelial cells. **A-I.** HDLEC were treated *in vitro* with conditioned media of NIH3T3 cells infected APLN lentivector. **A.** relative expression of APLN in NIH3T3 evaluated by RT-qPCR on NIH3T3 transduced by APLN lentivector. **B.** Expression of APLN in conditioned media evaluated by EIA. **C.** Representatives phospho-AKT/AKT and phosphor-Erk/Erk immunoblots of HDLEC treated with FBS, conditioned medium containing VEGF-C or APLN. **D.** Graphs represent quantification of phospho/total protein ratio of at least three independent experiments. All graphical data are mean ± s.e.m. *P < 0.05, twoway ANOVA. **E.** Representative images of scratch wound healing assay on HDLEC stimulated by VEGF-C or APLN. **F.** Quantification of migration (***p < 0.01). **G.** F-actin and VE-Cadherin immunostaining of HDLEC after APLN treatment reveals no effect on lymphatic endothelial monolayer junctions. **H.** Representatives phospho-eNOS/eNOS immunoblots of HDLEC treated with FBS, conditioned medium containing VEGF-C or APLN. **I.** Graphs represent quantification of phospho/total protein ratio of at least three independent experiments (*P < 0.05).

### APLN stimulates lymphatic collector pumping through eNOS activation

We next explored whether APLN could control vessel dilatation, in particular regarding collecting lymphatic vessels. APLN was described to activate eNOS phosphorylation in several cellular contexts and to promote blood vessel dilatation (Dray *et al*, 2008; Wysocka *et al*., 2018). We then investigated whether APLN was able to stimulate lymphatic collecting vessel dilatation and thus lymphatic pumping (Fig. 6A, supplemental movies 1-3). Lymph flow is in part driven in collecting lymphatics by autonomous contraction of smooth muscle cells. To evaluate the effect of APLN on collecting vessel contractions, we used intravital imaging method previously described (Liao *et al*, 2014)(Fig. 6A-C). Number of contractions and dilatation of vessels was assessed. Interestingly, we found that APLN stimulated lymphatic pumping by increasing the collector dilatation (Fig. 6A-B) without major effect on contraction frequency (Fig. 6C). This effect was completely reversed by the L-NAME, the nitric oxide synthase (NOS) inhibitor (Fig. 6A-C). Then, to evaluate the role of eNOS activation in response to APLN *in vivo* in LD context, LV-APLN treated mice were submitted to L-NAME treatment (Fig. 6D). Limb diameter was measured to assess the edema (Fig. 6E). Interestingly, L-NAME reversed the beneficial effect of APLN on LD confirming the lymphatic pumping as a major etiology of the pathology. We also observed an increase of edema two weeks after surgery in the presence of APLN + L-NAME (Fig. 6E). Lymphangiography revealed that L-NAME treatment also reverses APLN effect on lymphatic vascular network. Indeed, in APLN + L-NAME-treated mice, we observe pathological remodeling of lymphatic vessels with dermal backflow and abnormal lymphatic branching (Fig. 6F). Capillaries were also quantified using Lyve1 immunodetection on skin sections (Fig. 6G-I). An increase of lymphangiogenesis was observed systematically in LD limb, however we did not observe any differences between conditions (Fig. 6H). Nevertheless, regarding vessel area, APLN-treated mice displayed an increased dilatation that was inhibited by L-NAME treatment restoring a similar phenotype compared to control group (Fig. 6I). We also investigated fibrosis and surprisingly L-NAME had no effect on fibrosis (Fig. EV2). Taken together, our results show that APLN prevents LD by promoting collecting vessels pumping and remodelling of lymphatic vessels. This phenotype seems to be mediated in part by activating eNOS in lymphatic endothelial cells.

**Figure 6:**
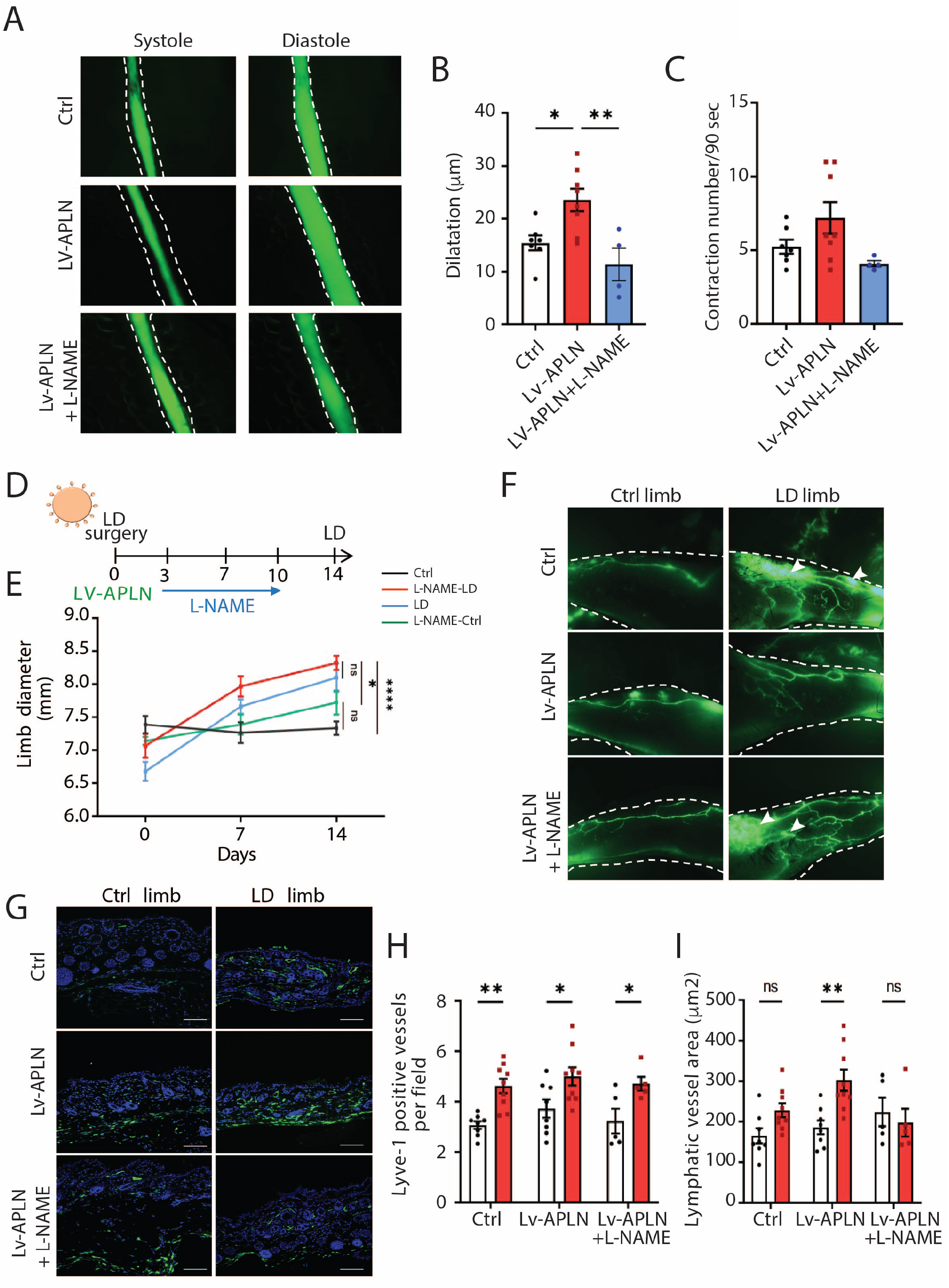
APLN-induced vasodilatation of lymphatic vessels is mediated by eNOS signalling. **A.** Contractile activity of collecting lymphatic vessels in control mice (n= 7) treated with APLN lentivector (n=8) and L-NAME (n=4) was investigated by filming autonomous collecting vessel contraction *in vivo*. **B-C.** graphs represent the number of vessels contraction per film (90 seconds) and the dilatation of collecting lymphatic vessels (differences between maximum and minimum diameter) (*p<0.05, **p < 0.01). **D.** Schematic of the experimental design of secondary LD mice model. **E.** Quantification of proximal limb swelling at 7 and 14 days after surgery on control limb, LD limb (*n=9*) or LD treated with APLN lentivector (*n=10*) followed or not by treatment with L-NAME (n=5) (*p<0.05, ****p < 0.001). **F.** Representatives images of lymphangiography from mice treated with LV-APLN and L-NAME. **G.** Skin sections were stained with Lyve 1 (green) to assess the number of lymphatic capillaries in control (n=9), LV-APLN (n=9) or LV-APLN + L-NAME (n=5) treated mice. **H-I.** Graphs show the number of lymphatic vessels (Lyve1^+^)(**H**) and the vasodilatation (vessel area) **(I)** according the experimental conditions (*p<0.05, **p < 0.01).

### APLN and VEGFC exhibit a synergistic effect on the regulation of gene expression related to collecting vessels maintenance

Most of the studies aiming at regenerating the lymphatic system have focused on the VEGF-C molecule. Nevertheless, VEGFC alone has appeared ineffective to improve lymphatic collector function in mice model of vascular injury, suggesting that it has to be combined with other molecules to fully restore the lymphatic drainage. Here we found that APLN controlled LD fibrosis, lymphatic function and contractility of collecting vessels. Our original approach aims at combining VEGF-C with APLN to obtain a synergistic effect for the treatment of secondary LD by targeting the entire lymphatic network from capillaries to collectors. When comparing gene expression profile of APLN-, VEGF-C or APLN+VEGF-C stimulated HDLEC, we observed similar induction of top 30 genes mostly related to extracellular matrix remodeling (Fig. 7). The majority (43) of the genes induced by VEGF-C are also induced by APLN (Fig. 7A-C)(Table S3). Half of the genes induced by the combination of APLN+VEGF-C are induced by APLN (Fig. 7D-F)(Table S4). When comparing induction of genes shared by the two molecules, the cooperative effect of APLN and VEGF-C remained focused on genes related to the microenvironmental maintenance of the lymphatic system (collagen 1A1 and 6A) and to the maturation of VEGF-C (ADAMTS2, E2F8)(Fig.7G). Also, 33 genes are upregulation by the combination of APLN and VEGF-C (Fig.7G). When comparing non treated-, VEGFC alone or APLN alone to APLN+VEGFC combination, we found an increase of genes necessary for collecting vessel function including connexin 37 and 47 (GJA4, GJC2) and claudin 5 (CDN5). The expression of genes related to VEGFC maturation was improved (ADAMTS2, CCBE1), whereas angiogenic genes were down-regulated (VEGFA, Flt1, KDR, KI67)(Fig.7H and I).

**Figure 7.**
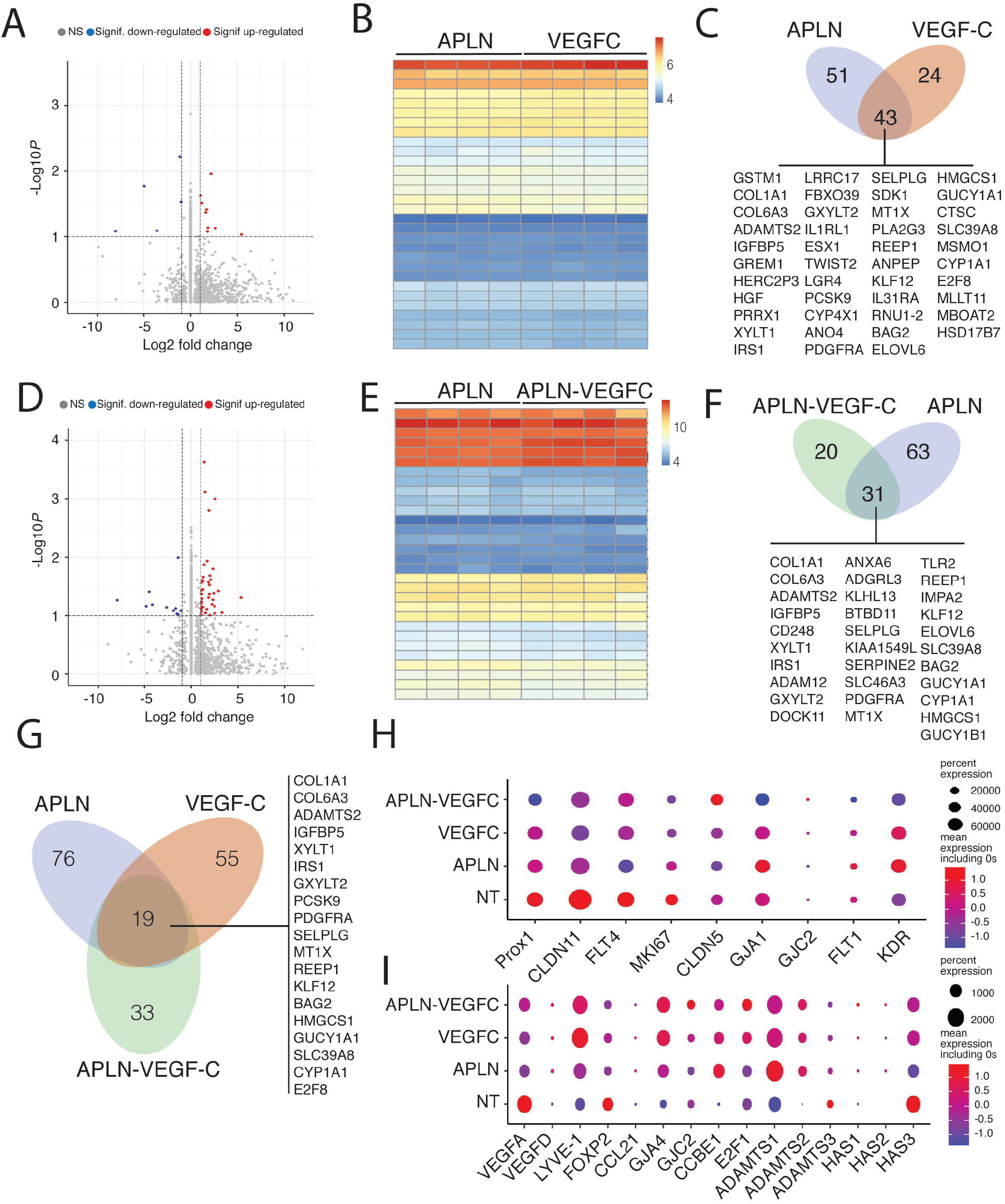
APLN and VEGFC exhibit complementary effect on lymphatic endothelial cells. **A-I.** Comparison of bulk RNA-sequencing in HDLEC treated with APLN-, VEGF-C-, or APLN +VEGF-C-conditioned media. **A-C.** Bulk RNA-sequencing in HDLEC treated with APLN- and VEGFC-conditioned medium. **A.** Volcano plot showing log2FC (Fold change) values calculated between APLN- and VEGFC-treated HDLECs. Red and blue dots: significantly (*p value adjusted* < 0.05) up- (log2FC > 0.5) and downregulated genes (log2FC < −0.5) respectively. **B.** Heat map comparison of the top 30 significantly regulated genes in APLN- and VEGFC-treated HDLEC. **C.** Schematic representation of the number of genes up regulated (43) by both APLN and VEGF-C. **DE.** Bulk RNA-sequencing in HDLEC treated with APLN and APLN-VEGFC-conditioned medium. **D.** Volcano plot showing log2FC (Fold change) values calculated between APLN- and APLN-VEGFC-treated HDLECs. Red and blue dots: significantly (*p value adjusted* < 0.05) up- (log2FC > 0.5) and downregulated genes (log2FC < −0.5) respectively. **E.** Heat map comparison of the top 30 significantly regulated genes in APLN- and APLN-VEGFC-treated HDLEC. **F.** Schematic representation of the number of genes up regulated (31) by both APLN and APLN-VEGF-C. **G.** Schematic representation of the number of genes up regulated (19) by both APLN, VEGF-C and APLN+VEGF-C. **H.** Dot plots showing the expression of known lymphatic markers in non treated (NT), APLN, VEGF-C and APLN+VEGF-C-treated HDLEC. **I.** Dot plots showing the expression of known lymphatic markers in NT, APLN, VEGF-C and APLN+VEGF-C-treated HDLEC.

### APLN-VEGFC RNA delivery: a new therapeutic option for secondary LD

In western countries, secondary LD develops mostly after cancer treatment, which makes ethical concern for the delivery of angiogenic molecules to cancer survival patients. For security reasons, we decided to use the next generation of vector called LentiFlash^®^ (Lf) which allows mRNA transient delivery from non-integrative viral particles. Thanks to its capacity for delivering several heterologous mRNA molecules, we then generated a Lf vector containing two mRNA sequences coding for VEGF-C and APLN to inject in the mouse model of LD (Fig. 8A). Lf efficiency is highly dependent of mRNA stability compared to lentivector that induces permanent expression of the transgene. We therefore first confirmed the presence of circulating APLN (Fig. 8B) and VEGFC (Fig. 8C) by ELISA, measurable 48 hrs after injection. We only observed a partial inhibition of limb swelling using APLN or VEGF-C mRNA alone (Fig. 8D and E). This could be expected, due to the restricted time of molecule expression. However, the APLN-VEGFC double mRNA Lf completely abolished limb swelling (Fig. 8F), reduced dermal backflow (Fig. 8G) and restored the lymphatic perfusion in the lymphedematous limb (Fig. 8G). This was associated with an increase in lymphatic vessel dilatation (Fig. 8H). Finally, to investigate whether Apelin-VEGFC mRNA could be a curative treatment for LD, we injected mice which developed LD (10 days after surgery)(Fig. 8I). In that context, Lf vector reversed LD swelling to go back to normal after 11 days. These data showed the synergistic effect of Apelin and VEGFC and demonstrated that this combination generates a significant therapeutic benefit despite the transient expression of the two transgenes, providing a perspective of LD treatment using “safe” RNA delivery vectors for patients who develop LD after cancer treatment.

**Figure 8.**
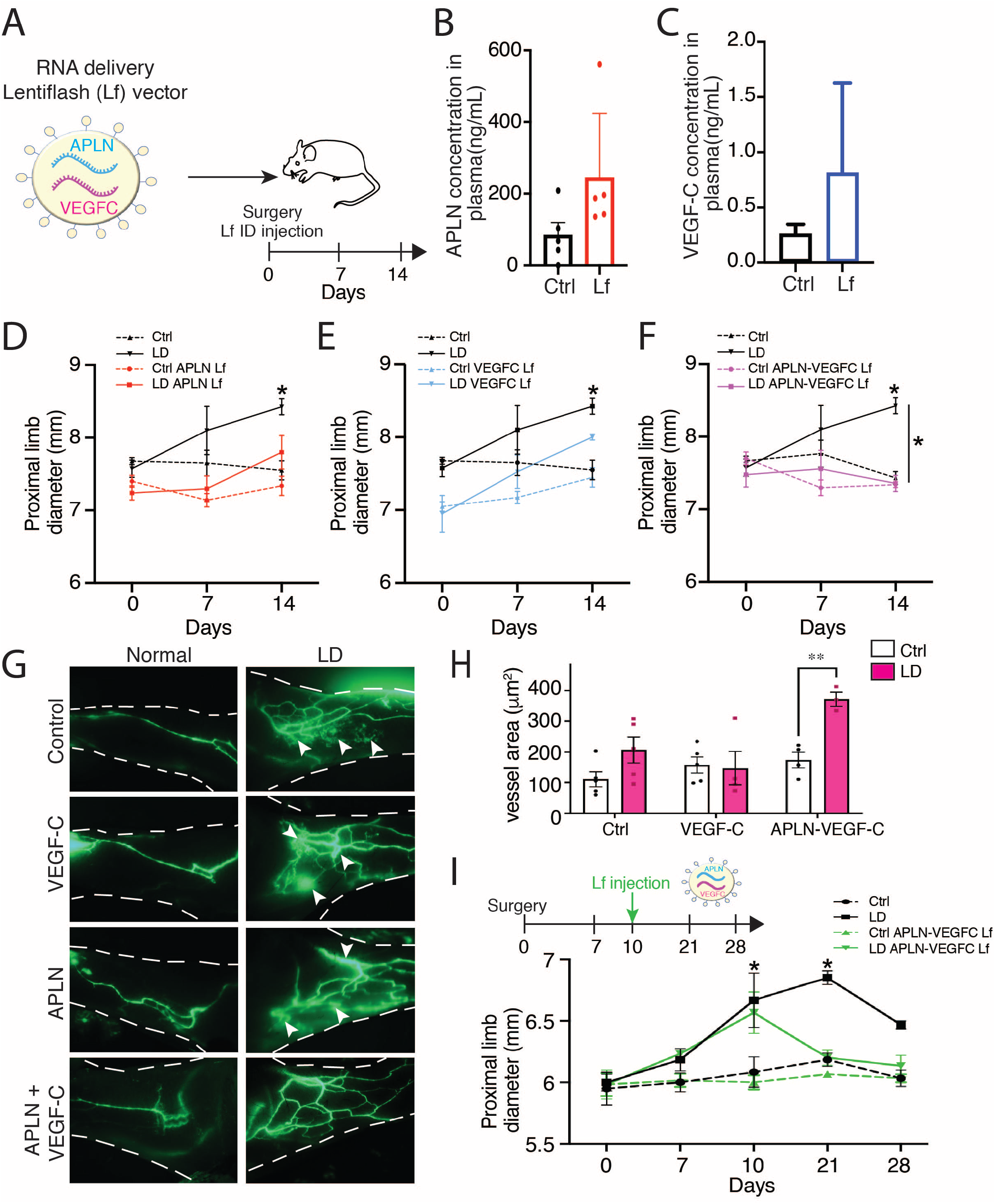
APLN-VEGF-C mRNA delivery: a new treatment option for LD. **A**. Schematic representation of the LentiFlash^®^ vector that encapsulates APLN and VEGF-C mRNA molecules for *in vivo* delivery. **B,C.** EIA dosage of circulating APLN (B) and VEGF-C (C) in plasma of control (n=5) or LentiFlash^®^-treated mice (n=5). **D.** Quantification of proximal limb swelling at 7 and 14 days after surgery on control limb, LD limb or LD treated with APLN LentiFlash^®^ vector (*n=10*) (*p<0.05). **E.** Quantification of proximal limb swelling at 7 and 14 days after surgery on control limb, LD limb or LD treated with VEGF-C LentiFlash^®^ vector (*n=10*)(*p<0.05). **F.** Quantification of proximal limb swelling at 7 and 14 days after surgery on control limb, LD limb or LD treated with APLN-VEGF-C LentiFlash^®^ vector (*n=10*)(*p<0.05). **G.** Representatives images of lymphangiography from mice treated with VEGF-C-, APLN-, or APLN-VEGF-C-LentiFlash^®^ vectors. **H.** Quantification of lymphatic dilatation in APLN-VEGF-C-treated mice (**p < 0.01). I. Quantification of proximal limb swelling in mice treated with with APLN-VEGF-C LentiFlash^®^ vector after LD development (10 days post-surgery) (*p<0.05).

## DISCUSSION

Despite large advances in the past decades for the understanding of the molecular mechanisms that drive the lymphatic function, LD, the most predominant pathology associated with lymphatic dysfunction remains an unmet medical need (Mercier *et al*, 2019). It is a chronic condition that affects millions of people worldwide. Many factors contribute to the etiology of the disease. Primary LD, an inherited disease, is induced by genetic mutation, whereas secondary LD occurs after cancer treatment or filarial infection (Mortimer & Rockson, 2014; Rockson, 2018). However, they all lead to comparable clinical signs: an accumulation of fluid and fat in the limb associated with fibrosis and hypervascularized dermis characterized by tortuous and leaky capillaries and hypoperfusion of deeper collecting vessels. Lymphoscintigraphies show a severe reduction of lymph node perfusion demonstrating that the lymphatic collecting vessels are still present, but cannot collect and drive the lymph properly. These observations support the regenerative therapeutic strategy aiming at combining molecules to 1/normalize the capillary territories and 2/ regenerate the lymphatic pumping in deeper adipose depots. Whereas it is now well established that VEGF-C, the major lymphangiogenic growth factor, is the best candidate the restore the lymphatic capillary network (Hartiala *et al*., 2020), its role on collecting remains less effective. Collecting vessels develop in an integrated adipose environment that is considerably modified during LD. In particular, adipose tissue synthesize many adipokines involved in the blood and lymphatic vessel integrity. It is therefore tempting to speculate that changes in adipokine production may affect the lymphatic collecting function. Among them, APLN has been described to be a key factor for stimulating LEC function (Kim *et al*., 2014). APLN is a bioactive peptide that induces signaling after binding to its G protein-coupled receptor APJ located at the surface of LEC. It stimulates lymphangiogenesis in cancer and participates to the restoration of precollecting lymphatics shape after myocardial infarction (Tatin *et al*., 2017). In addition to its effect on the endothelial monolayer, APLN is a robust antifibrotic molecule (Huang *et al*., 2016). Importantly, it was recently found to promote the non-sprouting expansion of vessels in intestinal crypt (Bernier-Latmani *et al*, 2022).

By performing gene expression analysis of dermolipectomies from women who developed secondary LD after breast cancer, we identified a significant decrease in APLN expression in LD. The crucial role of APLN in LD was confirmed in APLN KO mice that exhibit an aggravation of LD that can be rescued by an APLN-expressing lentivector. In the mouse model of LD, we identified that APLN improves LD condition by acting on two major hallmarks of the pathology: lymphatic function and tissue fibrosis. In LEC, APLN controls the expression of genes involved in extracellular matrix remodeling in line with its effect on tissue fibrosis. Interestingly, APLN also strongly stimulated the expression of CCBE1, a protein involved in the proteolytic activation of VEGF-C by ADAMTS3 (Jha *et al*., 2017). This could in part explain the increase of circulating VEGF-C concentration observed after APLN treatment. Importantly, in human, mutations in *CCBE1* were found to cause Hennekam syndrome, a congenital disease leading to LD, lymphangiectasia, and heart defects (Alders *et al*, 2013). Mechanistically, we found that CCBE1 gene expression is controlled by APLN that directly increase the fixation of E2F8, transcription factor on its promoter. Altogether, these data reinforce the APLN as a key factor in restoring the lymphatic function in LD.

Also, we found that the effect of APLN on the lymphatic collecting pumping was directly controlled by eNOS. NO production participates to the endothelial homeostasis by controlling the modulation of vascular tone as an adaptation of flow (Dimmeler *et al*, 1999). The endothelial NOS (eNOS) also regulates lymphatic homeostasis. In a mouse model of fibrosarcoma, eNOS mediates VEGF-C induced lymphangiogenesis and tumor lymphatic metastasis (Lahdenranta *et al*, 2009). Other studies have shown that eNOS affects the lymph flow via the collecting lymphatics, without affecting the diameter of capillaries (Hagendoorn *et al*, 2004). Also, NO bioavailability in pulmonary lymphatics was found to be impaired in limbs that exhibit chronically increased pulmonary blood and lymph flow (Datar *et al*, 2016). Here, we identified that the effect of APLN on the lymphatic collectors pumping is mediated by eNOS. APLN was previously described to modulates the aortic vascular tone by increasing the phosphorylation of Akt and eNOS in diabetic mice (Zhong *et al*, 2007). We found that the beneficial effect of APLN on lymphatic collecting vessels is mediated by this pathway suggesting that APLN can be at the origin of NO-mediated lymphatic pumping in many organs. The Akt-phosphorylation was found in a lesser extent than the phosphorylation induced by VEGF-C, however, it seems to be efficient to mediate its biological effects and suggests that APLN and VEGF-C have complementary effect to mediate biological actions. Therefore, we propose to evaluate the effect of APLN coupled to VEGF-C for the treatment of LD. However, an important ethical issue in treated cancer survivor patients is to reactivate the tumor with pro-lymphangiogenic therapy, even with more than five years without any recurrence. Therefore, the use of DNA delivery tools such as integrative lentiviral or AAV vectors rapidly appeared as not an optimal solution for treatment delivery due to uncontrolled protein expression duration and insertional mutagenesis risk. Another issue is the brief plasma APLN half-life which is less than five minutes (Japp & Newby, 2016). This could have been compensated by successive injections in the limb, however it would significantly improve the risk of infections and desmoplastic reaction, which are often seen in LD patients. We then decided to use a biological RNA delivery approach able to limit the dose requirement of a synthetic RNA-based therapeutic drug. LentiFlash^®^ technology based on a novel class of chimeric lentiviral platform allows the delivery of transient multiple biological mRNA molecules (Prel *et al*, 2015). LentiFlash^®^ is constructed using a bacteriophage coat protein and its cognate 19-nt stem loop, to replace the natural lentiviral Psi-mediated packaging system, in order to achieve active biological mRNA packaging into the lentiviral particles. Associated to a total removing of HIV sequences on the RNA packaged into lentiviral particle (LTR, PBS, RRE sequences), it enables the encapsidation of multiple and heterologous RNA sequences into the same LentifFlash particle without any risk of integration into the host genome. The present study shows that single APLN mRNA delivery mediated by LentiFlash^®^ exhibits less efficiency in reducing LD compared to integrative lentivector. However, when combined to VEGF-C, double mRNA delivery completely abolished LD and restored the lymphatic flow in the limb showing that mRNA delivery strategy allows enough synthesis of the two proteins to observe a beneficial effect.

In the past two years, we have seen the emergence of a novel class of mRNA vaccine, a highly efficient and low-toxic vectors. We believe that mRNA can treat many diseases including LD, a pathology with currently no treatments, in a different way than traditional medicine. The plasticity of LentiFlash^®^ allows the transient delivery of two different mRNA molecules, allowing here to stimulate the synergistic effect of APJ, a G protein-coupled receptor with VEGFR-3, a tyrosine kinase receptor. Based on the fact that LD remains a multifactorial pathology with lymphatic endothelial dysfunction, AT accumulation and fibrosis, we are convinced that multiple therapy will be the solution to cure this harmful condition. Therefore, we proposed to use the APLN-VEGF-C LentiFlash^®^ vector for a Phase I/II gene therapy clinical trial called Theralymph that is in the process of being launched in Toulouse University Hospital where our laboratory is located.

## Materials and Methods

### Human tissue specimen

Samples were obtained from archival paraffin blocks of 16 lipodermectomy specimens, obtained from patients with secondary LD, treated at Toulouse University Hospital, France, between 2015 and 2016. Samples were selected as coded specimens under a protocol approved by the INSERM Institutional Review Board (DC-2008-452), the Research State Department (Ministère de la recherche, ARS, CPP2, authorization AC-2008-452) and the Ethic Committee. When available, some control arm tissue samples removed for esthetical purpose in the same patients, were studied.

### Lymphofluoroscopy

Near-infrared fluorescence lymphatic imaging is used to visualize the initial and conducting lymphatics. 100 μg of indocyanine green (Pulsion^®^) was diluted in a volume of 0.5 mL of pure water before intradermal injection into the first interdigital space. Fluorescence imaging of the lymphatic flow was observed by fixing the camera (Photo Dynamic Eye, Hamamatsu^®^) 15 cm above the investigation field. Indocyanine green lymphography findings are classifiable into two patterns: normal linear pattern and abnormal dermal backflow pattern.

### Lymphoscintigraphy

Lymphoscintigraphy is used to diagnose the severity of lymphedema. It is a low radiation examination but forbidden during pregnancy and breastfeeding. Bilateral hypodermal injections are administered between the first and second fingers. The large size of 99m-technetium radiolabeled nanocolloid albumin is selectively entrapped by lymphatic capillaries and then drained by the lymphatic system. It enables comparative, functional and bilateral evaluation of the two upper limb inclunding axillary lymph-node uptake, lymphostasis, dermal back flow, and rerouting into the deep lymphatic system into the epitrochlear lymphnode.

### Mouse Model of LD

Mouse procedures were performed in accordance with EU and national regulation. C57Bl/6 mouse were provided by Envigo. All experiments have been approved by the local branch Inserm Rangueil-Purpan of the Midi-Pyrénées ethics committee. Secondary LD was established as previously described (Morfoisse *et al*., 2018). Briefly, LD was established in the left upper limb of 6 weeks-old C57Bl/6 female mice. A partial mastectomy of the second mammary gland is performed in association with axillary and brachial lymphadenectomy. Limb size was measured over time in the axillary region using calliper. Mice sustained edema for a period of 2-4 weeks. The day of the surgery, vehicle or APLN lentiviral vectors were injected intradermally in the LD limb (3 injections of 2 μL 10^6^TU/mL). For LentiFlash^®^ vector, vehicle, APLN, VEGFC, APLN-VEGFC Lf vectors were injected intradermally in the LD limb (200ng of p24 splited into 3 injections of 2 μL each). For L-NAME (N^G^-nitro-L-arginine methyl ester) (Sigma) treatment, L-NAME was resuspended in water (1 mg/ml) and mice were allowed to drink freely for 7 days.

### Lymphangiography

Two weeks after surgery, mice were anesthetized with an intraperitoneal injection of ketamine (100 mg/kg) (Zoletil 100, Virbac) and xylazine (10 mg/kg) (Rompun 2%, Bayer). FITC-Dextran (70 kDa, 2mg/mL, Sigma) was injected into the footpad of LD and control limb. The fluorescent molecule was taken up by the lymphatics and excluded from the blood vessels. After 5 min, the skin was analyzed under the modular stereo microscope discovery. V12 stereo (Zeiss).

### Histology

Skin of LD and control tissues from human and mice limb were embedded in paraffin and sectioned on a microtome. 5μm sections were cut and placed on Superfrost Plus slides. Tissue were deparaffinized, rehydrated and antigen unmasking was realized with pH 9 Tris solution (H-3301, Vector Laboratories) 5 min three times in a microwave. After cooling, slides were washed in PBS and then blocked with 5% BSA solution at room temperature in humid chamber. Sections were incubated with primary antibodies O/N at 4°C (Rabbit anti-human Lyve-1, Fitzgerald; Goat anti-murine Lyve1, R&D AF2125; rabbit anti-CD31, abcam Ab28364) and washed three times in PBS. Sections were incubated with the corresponding secondary antibodies conjugated to Alexa −488 or −594 for 1h at room temperature at 1/400 dilution. For fluorescence, nuclei were stained with DAPI. Images were acquired using inverted microscope (Leica, DMi8). Images were analysed with Fiji software.

### Evaluation of Fibrosis

Dermis fibrosis was evaluated with Masson’s trichrome coloration (MST-100T, Cliniscience). Skin sections were deparaffinized and tissue were stained according the manufacturer’s recommendations. Images were acquired on a nanozoomer slide scanner. Dermis size quantification was performed using at least 20 measurements of the length between epidermis and hypodermis per field.

### Collecting vessel contraction measures

Vessel contraction measurement of afferent collecting lymphatic vessels to the popliteal lymph node (PLN) were performed as described previously (Liao S. et al.). Briefly mice were anesthetized with an intraperitoneal injection of ketamine (100 mg/Kg) and xylazine (10 mg/kg). 6μL of FITC-Dextran was injected into the footpad of the lower right limb. The skin is carefully removed to expose afferent collecting lymphatic vessels to the PLN. Mouse is then placed into a petri dish and onto the stage of an inverted microscope (Leica, DMi8). 4 videos of 90 sec were acquired per mice and the number of contraction and dilatation (difference between maximum and minimum diameter) was analysed using Fiji software. To assess the effect of APLN, lentiviral vector was injected into the derma in the lower right limb 7 days before the experiment. For L-NAME study, mice were allowed to drink freely for 7 days.

### LentiFlash^®^ construction, production, purification, and quantitation by p24 ELISA assay

Four plasmids were used to produce recombinant LentiFlash^®^ particles in HEK293T cells: (i) the pLVGagPol plasmid encoding the viral gag and pol genes modified to harbor the PP7-Coat Protein (PCP) within the gag gene and referred to as pLF-GagPol ΔZF2_PCP (Mianne *et al*, 2022); (ii) the pVSVG plasmid encoding the VSV-G glycoprotein; and (iii) the two plasmids encoding each one RNA cargo, flanked by the PP7 bacteriophage aptamers to enable RNA mobilization into lentiviral particles through the interaction with the PP7 coat protein cloned in the Gag sequence‥ All newly generated constructs were verified by restriction enzyme digestion and sequencing. LentiFlash^®^ particles were produced in a 10-layer CellSTACK chamber (6360cm2, Corning) after transfection of the three plasmids in HEK293T cells using the standard calcium phosphate procedure. Twenty-four hrs post-transfection, the supernatant was discarded and replaced by fresh medium and cells were incubated at 37 °C in a humidified atmosphere of 5% CO_2_ in air. After medium change, supernatant was collected, clarified by centrifugation at 3000 g for 5 min, and microfiltered through 0.45-μm pore size sterile filter units (Stericup, Millipore). Supernatant was harvested several times, and finally, all samples were pooled (crude harvest). The crude harvest was concentrated and purified by ultrafiltration and diafiltration. For quantification, the p24 core antigen was detected directly in the viral supernatant with a HIV-1 p24 ELISA kit (Perkin Elmer), as specified by the supplier. The viral titer (expressed in physical particles per ml) was calculated from the p24 amount, knowing that 1 pg of p24 corresponds to 10E+4 physical particles.

### Enzyme Immuno Assay (EIA)

Concentration of APLN in the medium and in mouse plasma was determined using APLN EIA Kit (RAB0018, Sigma-aldrich) using the manufacturer’s recommendations. VEGFC ELISA kit was from R&D systems.

### Cell culture and treatments

Human dermal lymphatic endothelial cells (HDLEC) (single donor, juvenile foreskin, Promocell, C-12216, > 95% of the cells are CD31 positive and podoplanin positive) were cultured in Endothelial Cell Media MV2 (EGM-MV2, Promocell, C-22121). NIH3T3 were cultured in Dulbecco’s modified Eagle’s medium (DMEM, Sigma, D6429) supplemented with 10% fetal bovine serum (FBS, Gibco, 10270-06) and 1% penicillin-streptomycin. Endothelial cells were used at passage 3-6. Cells were cultured at 37°C in a 5% CO_2_ incubator. Culture medium was changed 3 times a week, and the cells were passaged 1/3. For collecting conditioned media, NIH3T3 were grown in 10 cm dishes, after reaching confluence, medium was removed and the cells were washed once with PBS. NIH3T3 were cultured overnight in 5 mL reduced serum medium optiMEM (Gibco), the medium was then collected and used for experiments. HDLEC were treated with 50% conditioned media/50% MV2-05% FBS.

### RNA extraction and Reverse transcriptase and qPCR

Total RNA was prepared using RNeasy kit (Qiagen 74106) according to the supplier’s instruction. 1 μg of RNA was retro-transcribed using high-Capacity cDNA Reverse Transcription Kit (Thermo Fisher Scientific, 4368813) containing Multi-Scribe Reverse Transcriptase according to the supplier’s instruction. Quantitative real-time PCR was performed using OneGreen FAST qPCR premix (Ozyme, OZYA008) on a StepOne Real-time PCR System (Thermofisher Scientific). All samples were analysed in duplicates. Data was normalized relative to HPRT mRNA levels. The list of primers is the following: THSB2: F-AGAGTCACTTCAGGGGTTTGC; R-TGGCAACCCTTCTTGCTTAGA, CCBE1: F-ACATGGTGAAAGCCGGAACT; R-TTGTTGGGGAGCAGAGCAAT, E2F8: F-CATGCTCGAGGACAGTGGTT; R-GCACTGCGTGAGAGGGATTA, APLN: F-GTTTGTGGAGTGCCACTG; R-CGAAGTTCTGGGCTTCAC, ADAMTS3: F-TTCCAGGAACCTCTGTTGCC; R-GCTGATCTCTTGTAGACAAC, FBN:F-ACCTCA ACAGATGGCTCTCG, R-GCAGCACTGCATTTT CGTCA, COL1A1: F-TGATGGGATTCCCTGGACCT; R-CCAGCCTCTCCATCTTTGC, HPRT: F-TGGCCATCTGCCTAGTAAAGC; R-GGACGCAGCAACTGACATTTC, E2F1: F-AGGAACCGCCGCCGTTGTTCCCGT; R-CTGCCTGCAAAGTCCCGGCCACTT, Leptin: F-GCTGTGCCCATCCAAAAAGTCC; R-CCCAGGAATGAAGTCCAAACCG, Adiponectin: F-GTGAGAAAGGAGATCCAGGTCTT; R-TTTCCTGCCTTGGATTCCCG

### Immunoblotting

Cells were scrapped and lysed in RIPA buffer (RIPA 2X, Biotech RB4476) supplemented with phosphatase inhibitor (PhosSTOP Easypack, Roche0490687001) and protease inhibitors (protease inhibitor cocktail, Sigma Aldrich). Lysates were centrifuged at 13500g for 10 minutes at 4°C. Supernatants were then collected and mix with Laemmli buffer containing dithiothreitol (DTT 1mM). Proteins were resolved on 4-15% SDS-PAGE gels and transferred to nitrocellulose membrane (Trans-Blot Turbo RTA transfer kit, #1704271, Biorad). Membranes were blocked for 1h at room temperature in 5%BSA-TBS-T (TBS-0.1%Tween 20) and probed with primary antibody overnight at 4°C. Antibody used are the following:

Phospho AKT: AKT-p ser473 (CS#4060S), AKT: santa cruz H136 (S8312), Phospho ERK: ERK1/2-p(MAPKp42/44) (Thr202/Thr204) (cell signalling #9106), ERK ERK1/2-(MAPKp42/44) (Thr202/Thr204) (cell signalling #9102), Phospho eNOS (cell signalling #9571S), eNOS (cell signalling #5880S).

After three washes in TBS-T, membranes were probed with HRP-conjugated secondary antibodies at 1/10000 dilution. Signals were visualized with chemiluminescence detection reagent (Sigma) on a Chemidoc (Biorad) digital acquisition system.

### Bulk RNA-Sequencing

#### RNA Sequencing on primary human lymphatic endothelial cells

Total RNA from HDLEC treated or not with conditioned media containing APLN were harvested 24h after treatment and isolated using the RNeasy mini kit (Qiagen). DNA digestion was performed using the RNase-Free DNase set (Qiagen). Total RNA was then subjected to ribosomal-RNA depleted RNA sequencing (RNA-Seq) protocol performed by Genewiz company using Illumina HiSeq, PE 2×150 configuration.

Sequence reads were trimmed to remove possible adapter sequences and nucleotides with poor quality using Trimmomatic v.0.36. Reads were then mapped to the Homo sapiens reference genome (GRCh38) using the STAR aligner v.2.5.2b. Gene hit counts were calculated by using feature Counts from the Subread package v. 1.5.2. The hit counts were summarized and reported using the gene_id feature in annotation file. Only unique reads that fell within exon regions were counted.

#### Differential gene expression analysis

After extraction of gene hit counts, the gene hit counts table was used for downstream differential expression analysis. Using DESeq2, a comparison of gene expression between APLN treated HDLEC against control HDLEC was performed. The Wald test was used to generate p-values and log2 fold changes. Genes with log2FC > 0.5 or log2FC < −0.5 and an adjusted p-value < 0.05 were defined as differentially expressed genes and used for the downstream analysis. The global transcriptional change across the two groups compared was visualized by a volcano plot.

#### Gene ontology (GO) analysis

A gene ontology analysis was performed separately on the statistically significant sets of upregulated and downregulated genes, using PANTHER software (version 16.0, http://pantherdb.org/). The Homo Sapiens reference list was used to cluster the set of significantly differentially expressed genes based on their biological processes or pathways and the overrepresentation of gene ontology terms was tested using Fisher exact test. All the GO terms with a False discovery rate (FDR) lower than 0.05 were considered significant and are listed in Supplementary Data.

#### Chromatin Immunoprecipitation (ChIP)

Human dermal lymphatic endothelial cells (HDLEC) non transduced or transduced with APLN Lentivector, were crosslinked for 15 min using 1% formaldehyde directly in the culture medium. 0.125 M of Glycine were then added for 5 min. After two washes with cold PBS, cells were scraped and frozen at −80°C. Cells were lysed to ChIP-IT Express Magnetic Chromatin Immunoprecipitation kit (Active Motif 53008). Optimal sonication conditions were determined previously in order to obtain DNA fragments of about 500 bp. Cells were sonicated in 350 μl final volume of Shearing buffer specific to the kit, using Diagenode Bioruptor Sonicator (7 cycles, 30 sec ON, 30 sec OFF in a water bath). DNA concentration was determined using a Nanodrop and 25 μg of chromatin were used by reaction. Experiments were then performed according to the manufacturer protocol. 4 μg of E2F8 antibody (Abcam, AB109596) were used by ChIP reaction. 10 μl of each sample were kept as Input. Reactions were incubated over night at 4°C. A mock sample without antibody was processed similarly. Prior to qPCR, DNA was purified using the Active Motif Chromatin IP DNA purification kit (58002), and eluted in 50 μl of DNase/RNase free water. 2 μl of purified chromatin was used for qPCR. In some experiments, ChIP reactions were supplemented with 10 ng of Drosophila melanogaster chromatin (spike in chromatin, Active motif, 08221011), and 1 μg of an antibody recognizing H2Av, a Drosophila specific histone variant, (spike in antibody, active motif, 61686), as an internal control for ChIP normalization. The primers used for qPCR are the following: CCBE1: F-CCTCCTCCGTTTTCTTGTT; R-TTGTCCTGAGCGGCTTTAAT, E2F1:F-AGGAACCGCCGCCGTTGTTCCCGT; R-TGCCTGCAAAGTCCCGGCCACTT.

### Cell Transduction

NIH3T3 were seeded in 6-well plates at 100 000 cells/well. After 24h, cells were transduced with 1 mL of APLN lentivector diluted in 1 mL of OptiMEM Media in the presence of proteamine sulfate at a final concentration of 5 μg/mL. The Control cells (NT) non transduced with the APLN-, VEGFC-, APLN+VEGFC-Lentivectors were processed at the same time and following the same protocol; in this case 2 mL of OptiMEM and 5 μg/mL of protamine sulphate were added to the cells. The media was replaced after 24h. Cells were grown to reach confluence, and then passed and amplified for further experiments. APLN transduction was verified by RT-qPCR.

### Immunoblotting

Cells were scrapped and lysed in RIPA buffer supplemented with phosphatase inhibitor and protease inhibitors. Lysates were centrifuged at 13500g for 10 minutes at 4°C. Supernatants were then collected and mix with Laemmli buffer containing dithiothreitol (DTT 1mM). Proteins were resolved on 4-15% SDS-PAGE gels and transferred to nitrocellulose membrane (Trans-Blot Turbo RTA transfer kit, #1704271, Biorad). Membranes were blocked for 1h at room temperature in 5% BSA-TBS-T (TBS-0.1%Tween 20) and probed with primary antibody overnight at 4°C. After three washes in TBS-T, membranes were probed with HRP-conjugated secondary antibodies at 1/10000 dilution. Signals were visualized with chemiluminescence detection reagent (Sigma) on a Chemidoc (Biorad) digital acquisition system. The primary antibodies used are the following: APLN (Genetex GTX37465), E2F8 ((Abcam, AB109596), Total Akt (Santa Cruz biotechnology sc-81434), Phosphorylated Akt (Cell Signaling 4060), total eNOS (Cell Signaling 9572), phosphorylated eNOS (Cell Signaling 9571), Actin (Santa Cruz biotechnology sc-47778).

### Statistical analysis

All results presented in this study are representative of at least three independent experiments. In all figures, ‘*n*’ represents the number of biological replicates. Data are shown as the mean ± standard error of the mean (s.e.m.). Statistical significance was determined by two-tailed Student’s *t* test, one-way ANOVA or two-way ANOVA test with Bonferroni post hoc test using Prism ver. 9.0 (GraphPad). Differences were considered statistically significant with a *P* value <0.05. symbols used are: ns > 0.05, * ≤0.05, **≤0.01, ***≤0.0001.

## AKNOWLEDGEMENTS

We thank the module of Clinical Investigation Center (CIC 1436) from Toulouse University Hospital (Toulouse, France), in particular Dr F. M. Lebrin for her outstanding assistance. We thank E Lhullier and C. Segura (GenoToul platform) for their technical support as well as M. Rousseau from the platform Anexplo Genotoul (Inserm US006, Toulouse, France) for their technical advices. We thank the imaging platform of I2MC institute (R. Flores). This work has received funding from the European Union’s Horizon 2020 research and innovation program named Theralymph under grant agreement no. 874708. This work has been supported by the Cancéropôle GSO, the Foundation for medical research (FRM), the Foundation ARC pour la Recherche contre le Cancer, the National Institute of Cancer (Inca).

## AUTHOR CONTRIBUTIONS

A. L. performed most of the experiments, analyzed the data, and contributed to experimental design. J.C. performed RNAseq experiment. E.B., F.P., M.N., F.M. contributed to the mouse experimental procedures and immunostaining. E.B. produced lentivectors. T.D.N. performed bioinformatical analysis. E.L., F.G., P.V., J.M.D., A.B.R., AC.P. contributed to the experimental design and to the analysis of the experiments. R.G., P.B. contributed to experimental design and production of the LentiFlash^®^ vector. B.C., J.MD., A.BR. contributed to the human tissue collection and analysis. B.G.S. designed the study, performed data analysis, and wrote the manuscript.

## THE PAPER EXPLAIN

### Problem

Secondary lymphedema (LD) is a chronic condition that affects millions of cancer survivors. It is the consequence of a severe lymphatic dysfunction leading to the accumulation of fluid and fibrotic adipose tissue (AT) in a limb. It is a characterized by tortuous lymphatic capillary network and a severe decrease in lymphatic collecting vessels pumping. Whereas the vascular endothelial growth factor-C (VEGF-C) is well known to promote lymphatic capillary growth, little is known about the regenerative control of the lymphatic collecting vessels.

### Results

Our study addresses the role of Apelin (APLN), a key bioactive peptide, in lymphatic endothelial cells (LECs) function. We identified the loss of APLN in human LD tissue biopsies. Our study shows that APLN knockout mice recapitulate the increase of edema associated with a dermal lymph backflow and skin fibrosis. In particular, lymphatic collector pumping largely depends on APLN-mediated eNOS pathway. Overall, our data shows that APLN restores collector function and participates to capillary homeostasis by stimulating VEGF-C maturation.

### Impact

Our study reveals that targeting APLN and VEGF-C might prove useful as novel therapeutical strategy to treat the entire limb lymphatic network in LD patients. Importantly, we propose to use safe and transient mRNA delivery strategy for patients who develop LD after cancer treatment.

---

**Table S1.** Gene expression changes in response to APLN treatment. Bulk RNAseq analysis of the upregulated genes.

| Gene,name | log2FoldChange | padj |
| --- | --- | --- |
| SELE | 2,812731725 | 6,48E-08 |
| MMP10 | 2,752225374 | 0,000369035 |
| HDAC9 | 2,741997597 | 0,004410612 |
| CRYAB | 2,635306806 | 0,000147694 |
| CLDN1 | 2,389292005 | 4,77E-09 |
| AC089983,1 | 2,36902799 | 0,000247378 |
| CCL5 | 2,276903012 | 0,000607478 |
| SERPINB2 | 2,270073695 | 0,010644328 |
| CXCL8 | 2,211382158 | 0,000798952 |
| CDCP1 | 2,201518538 | 0,000370379 |
| MYEOV | 2,18951514 | 0,007186895 |
| LTB | 2,040538173 | 0,008297478 |
| OIP5 | 1,941683429 | 0,003142472 |
| C15orf54 | 1,928461604 | 0,00857569 |
| CSF3 | 1,926259196 | 0,011845263 |
| S1PR3 | 1,904941869 | 2,01E-05 |
| CPA4 | 1,89809741 | 0,000731568 |
| PLAU | 1,896943904 | 5,38E-11 |
| NRG1 | 1,896230589 | 0,000443877 |
| FGF5 | 1,885713721 | 0,000792216 |
| ICAM1 | 1,818957189 | 0,008977731 |
| CX3CL1 | 1,818549284 | 0,013746926 |
| APLN | 1,810337986 | 9,10E-05 |
| MAP2 | 1,801050773 | 0,000359968 |
| VCAM1 | 1,783412264 | 7,09E-05 |
| IL7R | 1,776145301 | 5,84E-05 |
| IL32 | 1,765738581 | 1,52E-05 |
| RGS7 | 1,750200793 | 0,03119854 |
| SERPINE1 | 1,733383007 | 2,27E-05 |
| AL590004,4 | 1,724948793 | 0,014089116 |
| AL121718,1 | 1,710482389 | 0,02231266 |
| KRT7 | 1,707903162 | 3,03E-47 |
| LINC01013 | 1,700691575 | 0,027792639 |
| LINC02407 | 1,69104104 | 0,015865541 |
| CD34 | 1,668560141 | 0,005604739 |
| ENOX1 | 1,668378107 | 1,60E-07 |
| AC009549,1 | 1,665659241 | 0,000766355 |
| RGS4 | 1,657093433 | 7,05E-19 |
| TRNP1 | 1,633496667 | 4,68E-16 |

**Table S2.** Gene expression changes in response to APLN treatment. Bulk RNAseq analysis of the downregulated genes.

| Gene,name | log2FoldChange | padj |
| --- | --- | --- |
| NTN1 | -2,0426424 | 9,54E-20 |
| GPR183 | -2,0409517 | 0,0151189 |
| MATN2 | -1,9126348 | 4,06E-15 |
| ZNF853 | -1,8193332 | 1,58E-06 |
| ADAMTS12 | -1,7470107 | 0,01345412 |
| ITGA8 | -1,7214137 | 0,00840882 |
| ADAMTS15 | -1,6507767 | 0,01705855 |
| STAB2 | -1,6417797 | 8,22E-10 |
| PCDH17 | -1,6337697 | 7,96E-06 |
| ABCA8 | -1,5882342 | 0,01187511 |
| RASSF10 | -1,5692227 | 2,79E-11 |
| ELN | -1,5485884 | 0,00206406 |
| RASL10A | -1,5108805 | 0,00368074 |
| GRIK1 | -1,4859747 | 1,89E-06 |
| AQP1 | -1,4781154 | 0,00578527 |
| FZD10-AS1 | -1,4640296 | 0,00338056 |
| STXBP6 | -1,4485271 | 3,83E-05 |
| ABCG2 | -1,3954647 | 0,00547247 |
| ZDHHC8P1 | -1,3727316 | 0,0442768 |
| F8 | -1,3398527 | 5,65E-09 |
| OXTR | -1,2639102 | 4,46E-07 |
| SCUBE3 | -1,2587987 | 4,39E-10 |
| SGSM1 | -1,2151541 | 7,92E-09 |
| TRMT9B | -1,1971681 | 0,00277753 |
| AC068580,1 | -1,1711234 | 0,00263639 |
| ANGPTL4 | -1,1455021 | 7,96E-09 |
| CLEC10A | -1,1383706 | 0,03671307 |
| LGR4 | -1,1359248 | 0,02576944 |
| IL33 | -1,1323922 | 0,00019155 |
| C2CD4C | -1,1260407 | 0,00139968 |
| CPNE5 | -1,1195582 | 2,82E-07 |
| FZD10 | -1,119268 | 2,69E-07 |
| ALDH5A1 | -1,1190964 | 0,01728336 |
| ADD3-AS1 | -1,1151456 | 3,28E-06 |
| LAMC3 | -1,1089873 | 6,10E-07 |
| SIPA1L2 | -1,1009693 | 0,00036695 |
| C10orf128 | -1,0985672 | 0,01944283 |
| IGF2 | -1,0928267 | 0,00339568 |
| REEP1 | -1,0831336 | 4,53E-08 |

**Table S3.** Gene expression changes in response to VEGFC treatment. Bulk RNAseq analysis of the upregulated genes.

| Gene,name | log2FoldChange | pvalue |
| --- | --- | --- |
| GSTM1 | 24,6440679 | 2,09E-11 |
| COL3A1 | 12,5362136 | 2,84E-06 |
| COL1A1 | 10,1028955 | 3,00E-06 |
| COL6A3 | 9,35723896 | 1,22E-05 |
| ADAMTS2 | 7,82327731 | 0,00017456 |
| IGFBP5 | 7,65579404 | 5,01E-05 |
| GREM1 | 7,34519332 | 4,68E-05 |
| AF165147,1 | 5,94252292 | 2,60E-05 |
| NTM | 5,8860802 | 3,00E-05 |
| HERC2P3 | 5,82767126 | 0,0003209 |
| HGF | 5,66483328 | 0,00017597 |
| PRRX1 | 4,7327915 | 0,00015021 |
| XYLT1 | 4,59041849 | 0,00021529 |
| IRS1 | 4,19232602 | 1,12E-05 |
| LRRC17 | 4,10894287 | 0,00015412 |
| GPAT2P1 | 3,97643673 | 6,96E-05 |
| FBXO39 | 3,75275673 | 0,00024014 |
| GXYLT2 | 3,59872809 | 0,00012651 |
| IL1RL1 | 3,23930623 | 0,00033524 |
| ESX1 | 2,8650003 | 0,00017638 |
| TWIST2 | 2,74510441 | 0,00015814 |
| LGR4 | 2,58878342 | 0,00016059 |
| PCSK9 | 2,58105772 | 7,42E-07 |
| CYP4X1 | 2,50782533 | 0,0002746 |
| ANO4 | 2,49821267 | 0,00024518 |
| HIST1H1A | 2,34992625 | 9,54E-11 |
| PDGFRA | 2,33585253 | 0,00010922 |
| HIST1H3J | 2,33141774 | 9,00E-12 |
| SELPLG | 2,24774514 | 3,66E-05 |
| SDK1 | 2,23247851 | 0,00033843 |
| HIST2H2BF | 2,11705463 | 1,76E-07 |
| MT1X | 2,00950211 | 5,24E-10 |
| PLA2G3 | 2,00032749 | 2,82E-05 |
| HIST1H2BL | 1,96650297 | 7,19E-07 |
| REEP1 | 1,80691562 | 1,52E-06 |
| HIST2H3D | 1,7979865 | 0,00016134 |
| ANPEP | 1,75818463 | 0,00031543 |
| KLF12 | 1,69231642 | 7,55E-05 |
| AK7 | 1,63217642 | 0,00016331 |
| IL31RA | 1,60992486 | 2,79E-05 |
| RNU1-2 | 1,47684928 | 1,61E-05 |
| BAG2 | 1,47215967 | 5,97E-05 |

**Table S4.** Gene expression changes in response to APLN-VEGFC treatment. Bulk RNAseq analysis of the upregulated genes.

| Gene,name | log2FoldChange | padj |
| --- | --- | --- |
| COL3A1 | 13,51297701 | 0,000494047 |
| COL1A1 | 11,04361238 | 0,000415896 |
| COL6A3 | 9,413802034 | 0,00470325 |
| ADAMTS2 | 8,27465003 | 0,034461364 |
| IGFBP5 | 7,452186173 | 0,008385024 |
| CD248 | 5,893908153 | 0,033065433 |
| TMEM200A | 5,861334143 | 0,009638981 |
| NTM | 5,5709033 | 0,023355587 |
| CARMN | 4,915481566 | 0,008503535 |
| XYLT1 | 4,510397241 | 0,00054922 |
| IRS1 | 4,395069034 | 0,007227541 |
| GPAT2P1 | 4,079233898 | 0,047483092 |
| ADAM12 | 3,908667028 | 0,000949715 |
| GXYLT2 | 3,329572904 | 0,046590834 |
| DOCK11 | 3,144087614 | 0,039312035 |
| ANXA6 | 3,026947231 | 0,007227541 |
| ADGRL3 | 2,897773947 | 0,023355587 |
| PCSK9 | 2,804968682 | 0,002364173 |
| KLHL13 | 2,75082358 | 0,036755614 |
| HIST1H3J | 2,548542987 | 1,35E-05 |
| HIST1H1A | 2,494982926 | 7,71E-06 |
| BTBD11 | 2,419793652 | 0,039530643 |
| SELPLG | 2,40537126 | 0,009804674 |
| KIAA1549L | 2,391485078 | 0,039186999 |
| SERPINE2 | 2,239191837 | 0,049204756 |
| SLC46A3 | 2,22965513 | 0,032557454 |
| HIST2H2BF | 2,069223465 | 0,034938273 |
| HIST2H3D | 2,053137471 | 0,010150563 |
| PDGFRA | 2,022277061 | 0,041201923 |
| HIST1H2BL | 1,995288276 | 0,032766325 |
| SSUH2 | 1,90226196 | 0,015561785 |
| MT1X | 1,796160767 | 0,000834822 |
| TLR2 | 1,713696714 | 0,032557454 |
| REEP1 | 1,702904635 | 0,000787739 |
| HIST1H3D | 1,645591486 | 0,000891381 |
| IMPA2 | 1,623196651 | 0,038381511 |
| NMRAL2P | 1,547984923 | 0,004786025 |
| KLF12 | 1,527556133 | 0,047483092 |
| ELOVL6 | 1,464493319 | 0,002051263 |

## EXPANDED VIEW FIGURE LEGEND

**Figure EV1. Effect of APLN vector on skin angiogenesis.**

**A.** Skin sections of control or LD limb of control or LV-APLN treated mice were stained for CD31 (red) DNA was stained with DAPI. Scale bar = 100μm. **B.** Quantification of blood vessel per field according the limb and treatment of mice (*n=9* control and *n=8* APLN treated mice). All graphical data are mean ± s.e.m.

**Figure EV2. Effect of L-NAME on dermis fibrosis.**

**A.** Fibrosis was evaluated by Masson’s trichrome staining of the skin in control mice (*n= 9*) treated with APLN lentivector (*n=9*) and LV-APLN + L-NAME (n=5). **B.** graph display the quantification of dermis thickness. All graphical data are mean ± s.e.m, **P < 0.01.

**Expanded Video 1. Limb collecting vessel contraction in WT mice.**

Vessel contraction measurement of afferent collecting lymphatic vessels to the popliteal lymph node.

**Expanded Video 2. Limb collecting vessel contraction in APLN-treated mice.**

Vessel contraction measurement of afferent collecting lymphatic vessels to the popliteal lymph node from mice treated with APLN lentiviral vector injected intreadermally in the lower right limb 7 days before the experiment.

**Expanded Video 3. Limb collecting vessel contraction in APLN + L-NAME treated mice.**

Vessel contraction measurement of afferent collecting lymphatic vessels to the popliteal lymph node from mice treated with APLN lentiviral vector and L-NAME 7 days before the experiment.

